# Increased expression of genetically-regulated *FLT3* implicated in Tourette’s Syndrome

**DOI:** 10.1101/812420

**Authors:** Calwing Liao, Veikko Vuokila, Hélène Catoire, Fulya Akçimen, Jay P. Ross, Cynthia V. Bourassa, Patrick A. Dion, Inge A. Meijer, Guy A. Rouleau

## Abstract

Tourette’s Syndrome (TS) is a neurodevelopmental disorder that is characterized by motor and phonic tics. A recent TS genome-wide association study (GWAS) identified a genome-wide significant locus. However, determining the biological mechanism of GWAS signals remains difficult. Here, we conduct a TS transcriptome-wide association study (TWAS) consisting of 4,819 cases and 9,488 controls and found that increased expression of *FLT3* in the dorsolateral prefrontal cortex (DLPFC) is associated with TS. We further showed that there is global dysregulation of *FLT3* across several brain regions and probabilistic causal fine-mapping of the TWAS signal prioritizes *FLT3* with a posterior inclusion probability of 0.849. We validated the gene’s expression in 100 lymphoblastoid cell lines, establishing that TS cells had a 1.72 increased fold change compared to controls. A phenome-wide association study points towards *FLT3* having links with immune-related pathways such as monocyte count. We also identify several splicing events in *MPHOSPH9, CSGALNACT2* and *FIP1L1* associated with TS, which are also implicated in immune function. This analysis of expression and splicing begins to explore the biology of TS GWAS signals.

## Introduction

Tourette’s Syndrome (TS) is a neuropsychiatric disorder that is characterized by motor and phonic tics^1^. The onset of the disorder is typically between the age of 5-7 years. TS has been shown to have a large genetic component, in which the first-degree relatives of TS patients have a 10- to 100-fold higher rate of TS compared to the general population^2,3^. Past genetic studies of TS have identified several implicated genes such as *CELSR3*, a gene where recurrent do novo variants are found in probands^4^. Furthermore, a recent genome-wide association study (GWAS) identified a genome-wide significant hit on chromosome 13, rs2504235, which is within the *FLT3* (*Fms Related Tyrosine Kinase 3*) gene^5^. Although GWAS is a powerful method for identifying associated genetic loci, it is often difficult to interpret the biological effects of significant hits.

Recently, a transcriptomic imputation (TI) methodology was developed to allow for the integration of genetic and expression data from datasets such as the CommonMind (CMC) and Genotype-Tissue Expression (GTEx) consortia^6,7^. The derivation of panels involves a machine-learning approach to characterize the relationship between gene expression and genotypes, making tissue-specific predictive models. The TI methodology can leverage these reference imputation panels from these consortia and identifies the genetic correlation between imputed expression and GWAS data^6^. Ultimately, TI allows for better characterization of GWAS data by prioritizing tissue-specific genes associated with disease^8^. This methodology has already been used to prioritize genes in many different traits^9–11^.

To identify genetically regulated genes associated with TS, we conducted a transcriptome-wide association study (TWAS) of the current largest TS cohort of 4,819 cases and 9,488 controls^5^. Brain-specific panels were derived from the CMC and GTEx 53 v7. The TWAS revealed the expression of *FLT3* to be increased across many brain tissues in TS, with the largest effect in the dorsolateral prefrontal cortex (DLPFC). This is consistent with previous studies that also implicated the DLPFC region in TS. Given that *FLT3* is expressed in lymphoblasts, we additionally measured the RNA expression of *FLT3* in 100 lymphoblastoid cell lines (LCL;50 cases and 50 controls). There was an increased expression in *FLT3* in LCL derived from TS cases, consistent with TWAS results. The top hit for the omnibus test also identified *ATP6V0A2* as a putative gene associated with TS, however, this was not genome-wide significant. In conclusion, increased expression of *FLT3* was implicated through TWAS across several brain tissues and expression in lymphoblastoid cell lines.

## Results

### Transcriptome-wide significant hits

To identify genes associated with TS, a TWAS was conducted using FUSION. The strongest significant hit was *FLT3*, with increased expression (Z=4.67, P=2.98E-06) in the DLPFC (Table 1). Interestingly, the gene also had increased expression in the brain cortex, hippocampus, anterior cingulate cortex, frontal cortex, cerebellum, and cerebellar hemispheres suggesting a global dysregulation across brain tissue types. The gene *DHRS11* was also implicated (Z=4.26, P=2.01E-05), although not genome-wide significant. An omnibus test also identified the top two genes: *ATP6V0A2* (P=3.70E-05) and *NEB* (P=1.72E-04) (Table 1).

**Table 1.** TWAS genes with association to Tourette’s Syndrome

| Gene | Method | Tissue | P-value | Permutation<br>P-Value | Z-score |
| --- | --- | --- | --- | --- | --- |
| *FLT3* | Expression | Dorsolateral prefrontal cortex | 2.98E-06 | 0.00437 | 4.67226 |
| *FLT3* | Expression | Cortex | 3.24E-06 | 0.01289 | 4.6551 |
| *FLT3* | Expression | Hippocampus | 7.87E-06 | 0.0108 | 4.46875 |
| *FLT3* | Expression | Anterior cingulate cortex BA24 | 8.12E-06 | 0.007 | 4.462 |
| *FLT3* | Expression | Frontal cortex BA9 | 8.30E-06 | 0.0185 | 4.4574 |
| *FLT3* | Expression | Cerebellum | 1.44E-05 | 0.011 | 4.3382 |
| *DHRS11* | Expression | Substantia nigra | 2.01E-05 | 0.0153 | 4.26398 |
| *FLT3* | Expression | Cerebellar hemisphere | 2.43E-05 | 0.0096 | 4.22163 |
| *ATP6V0A2* | Omnibus | - | 3.70E-05<br>(Nominally significant) | - | - |
| *NEB* | Omnibus | - | 0.000172<br>(Nominally significant) | - | - |
| *MPHOSPH9* | Splicing | Dorsolateral prefrontal cortex | 1.58E-05 | 0.01509 | -<br>4.317536 |
| *FIP1L1* | Splicing | Dorsolateral prefrontal cortex | 2.55E-05 | 0.00184 | 4.21065 |
| *CSGALNACT2* | Splicing | Dorsolateral prefrontal cortex | 3.39E-05 | 0.00090 | 4.14582 |

### Splicing in Tourette’s Syndrome

To identify splicing events associated with TS, a TWAS was done using CMC splicing data. There were no significant hits, however, there were several significant genes after permutation. The top three hits were *MPHOSPH9* (Z=-4.32, P=1.58E-05), *FIP1L1* (Z=4.21, P= 2.55E-05) and *CSGALNACT2* (Z=4.14, P=3.39E-05) (Table 1). However, we also caution on the interpretability of the effect direction given that alternatively spliced exons are typically negatively correlated^9^.

### Fine-mapping of *FLT3* locus

To determine whether *FLT3* is the putatively causal gene on the DLPFC, FOCUS was used to assign a probabilistic inclusion probability for genes at the TWAS region^12^. For the region 13:27284583-13:29257379 (hg19 coordinates), the *FLT3* gene had the highest posterior inclusion probability (PIP) of 0.849 and was included in the 90% credible gene set (Figure 1).

**Figure 1.**
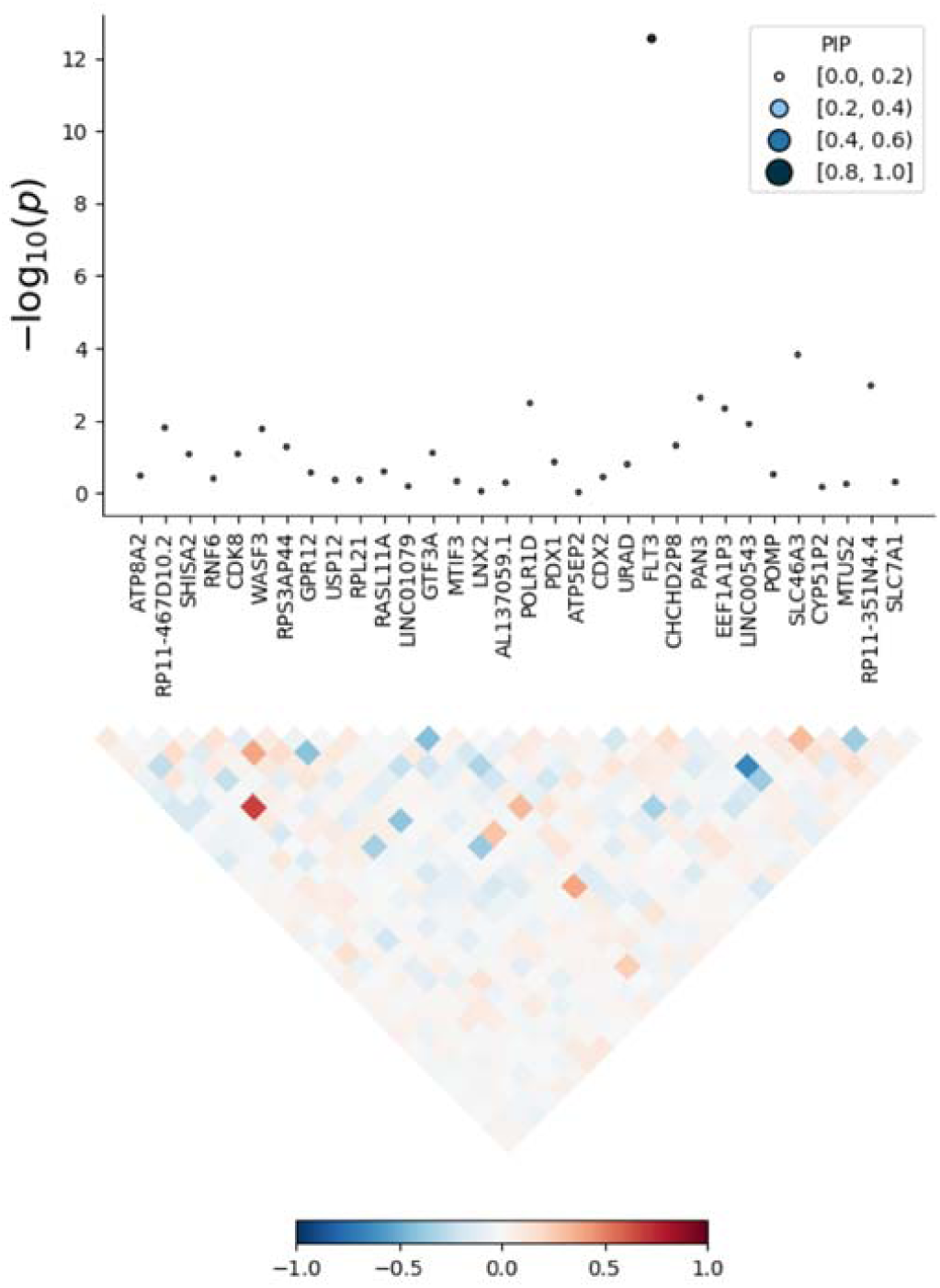
Fine mapping of chromosome 13 TWAS signal. PIP is the posterior inclusion probability. TWAS p-values derived from FOCUS are on the Y-axis and genes within the locus are on the X-axis. The local correlation structure is shown in the bottom half of the figure.

### RT-qPCR of *FLT3*

To test normality of qPCR data, a Shapiro-Wilk test was done. It was found that the ΔCT values (a measure of expression) were normally distributed (W=0.99, P=0.70). Next, an ANOVA was done and determined that the disease status explained a significant proportion of variance in the dataset (F=7.06, P=0.0095). A Tukey test showed that TS patients had significantly higher expression of *FLT3* compared to controls, with a ΔCT difference 0.780 (P=0.009) (Figure 2). The corresponding fold change is +1.72 higher in TS than controls. The effect size was determined to be moderate-large (Cohen’s F, 0.30).

**Figure 2.**
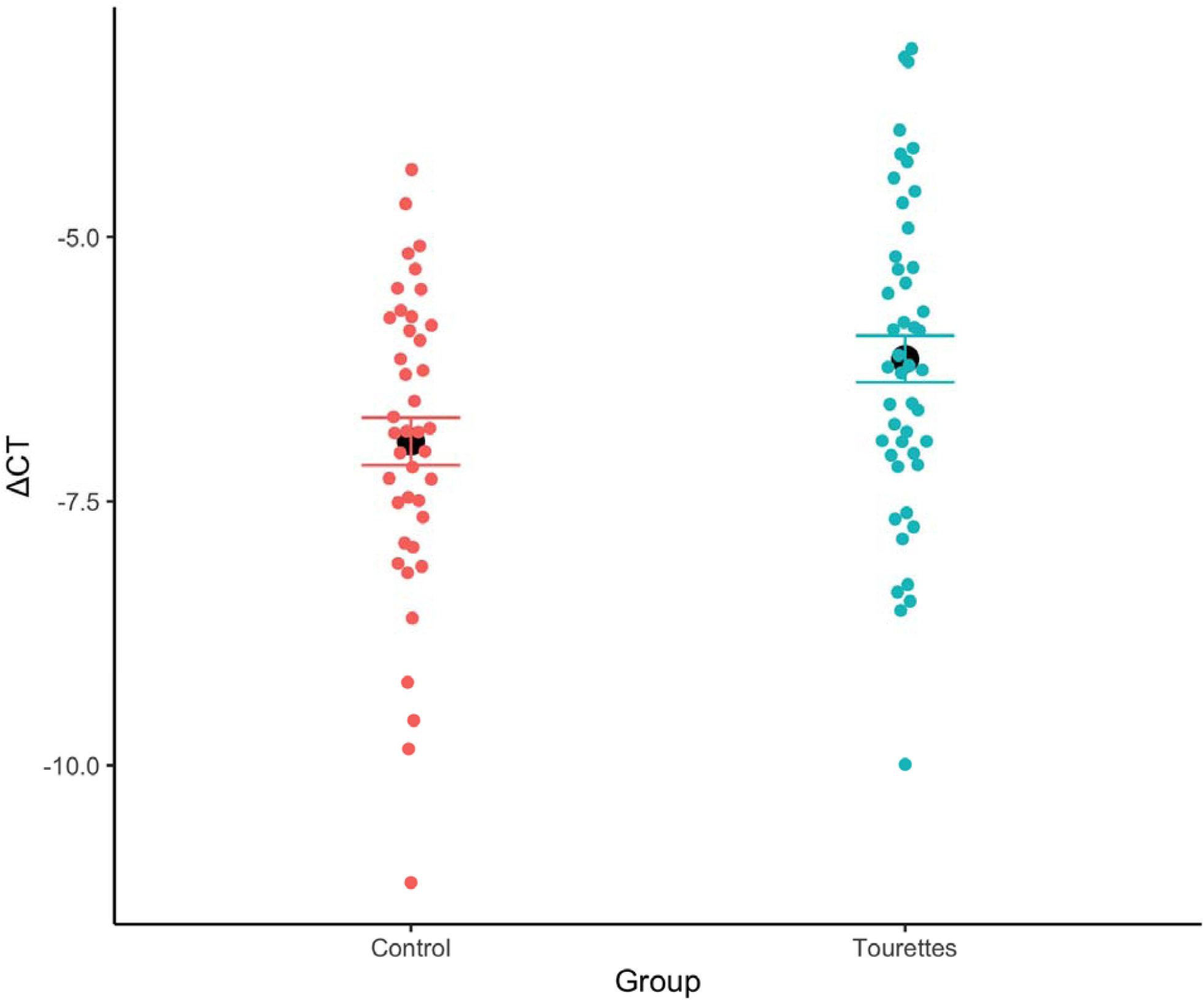
RNA Expression differences between Tourette’s Syndrome patients and controls in lymphoblastoid cell lines. Black dot represents the mean of the data and error bars are ±SE.

### Phenome-wide association study of *FLT3*

To identify phenotypes associated with the *FLT3* gene, a regional phenome-wide association study (pheWAS) was done. The pheWAS identified several immunological traits associated with *FLT3* such as monocyte count (3.87E-40) and percentage of white blood cells (1.42E-21) (Figure 3).

**Figure 3.**
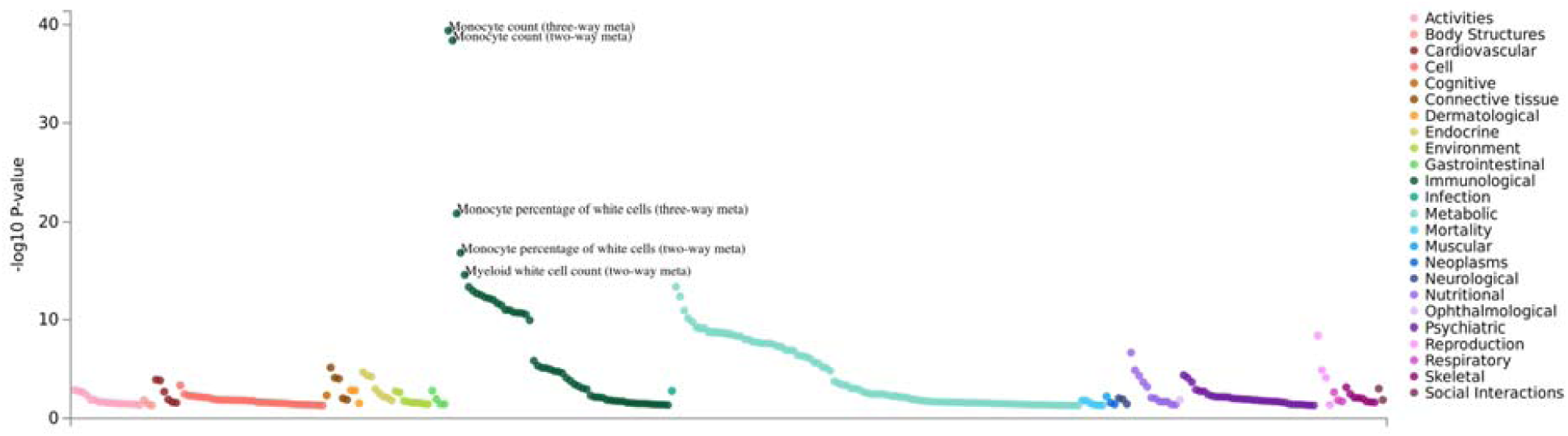
Regional phenome-wide association study of *FLT3*. Colours indicate respective domains.

## Discussion

While recent GWAS for TS have successfully identified risk loci, the biological relevance of these associations remains unknown. Here, we conduct a TWAS using the summary statistics of over 14,000 individuals from the most recent TS GWAS^5^. This approach allows for imputation of expression by leveraging genotype-expression reference panels. From this, we identified an increased *FLT3* expression as the top hit in the DLPFC and additionally found an increase in expression across most brain tissue types, suggesting global dysregulation. Validation of expression in LCL prepared from TS cases found an increased or an increase in RNA expression compared to LCL from control individuals. The *FLT3* gene encodes for a tyrosine-protein kinase and it has been associated with inflammation and immune function^13,14^. This could point towards immunity in TS as a putative biological mechanism. Furthermore, the pheWAS identified that *FLT3* is associated with immunological traits such as monocyte count and white blood cell counts. Interestingly, previous studies have demonstrated that TS patients have significantly higher levels of monocytes compared to healthy controls^15^. This could suggest dysregulation of monocytes partially due to increased expression of *FLT3*, which may contribute towards pathogenicity of TS. Furthermore, fine-mapping the TWAS hit demonstrated that *FLT3* was in the 90% causal credible-set with a PIP of 0.849 for the DLPRC. This further stipulates that *FLT3* is the correctly prioritized gene at this locus.

The splicing TWAS identified several putatively associated genes (*MPHOSPH9, FIP1L1*, and *CSGALNACT2*) associated with TS, suggesting that both splicing and genetically regulated genes are implicated in TS. The *MPHOSPH9* gene encodes for a protein that regulates cell cycling^16^. This gene has been implicated in multiple sclerosis, which is an inflammatory disease of the central nervous system^16^. Additionally, *FIP1L1* is associated with pre-mRNA 3’-end formation and has been implicated in immunological function by cooperating with IL-5^17,18^. These findings further support the hypothesis that the pathophysiology of TS may include or involve the immune system. Understanding the role of immunity in TS may elucidate the link between streptococcal infections and tic exacerbations as proposed in the pediatric autoimmune neuropsychiatric disorders associated with streptococcal infections (PANDAS) hypothesis^1,19^. The *CSGALNACT2* gene encodes for chondroitin sulfate protein, which is involved in the brain matrix^20^. A previous meta-analysis of ADHD and TS showed implication of sulfuration of chondroitin, suggesting potential relevance^21^.

We conclude this study with some caveats and potential future directions. First, TWAS signals can putatively be confounded due to expression imputation from weighted linear combinations of SNPs. Because of this, some of these SNPs may be associated with non-regulatory mechanisms that inflate the test statistic. A second caveat is that there is currently no available replication cohort, given that the largest GWAS for TS was used for this study. Future work could look at integrating single-cell sequencing data with TS GWAS to determine single-cell cis-eQTL regulated genes. Furthermore, individual TWAS risk could be investigated in independent cohorts. A third caveat is that a given gene may be influenced by genetic regulators independent of cis-eQTLs and sQTLs but still have downstream effect on TS. In conclusion, we identify the *FLT3* gene as likely involved in TS with increased expression found by TWAS and in lymphoblastoid cell lines of patients. We further identify several significant genes associated with aberrant splicing and point towards immunity in the pathogenesis of TS.

## Methods

### Genotyping data

Public summary statistics were obtained from the Psychiatric Genetics Consortium through the OCD & Tourette Syndrome group. Details on the participant ascertainment and quality control steps were previously described in the 2019 TS GWAS^5^.

### Transcriptomic imputation

Imputation was done by using reference panels from FUSION that were derived from consortia datasets of tissue-specific gene expression integrated with genotypic data. The CommonMind Consortium (CMC) and brain tissue panels from GTEx 53 v7 were used. Genes with a strict permutation empirical p-value <0.05 were considered significant. FUSION was used to conduct the transcriptome-wide association testing. The 1000 Genomes v3 LD panel was used for the TWAS. FUSION utilizes several penalized linear models, such as GBLUP, LASSO, Elastic Net. Additionally, a Bayesian sparse linear mixed model (BSLMM) is used. FUSION computes an out-sample R^2^ to determine the best model by performing a fivefold cross-validating of every model. After, a multiple degree-of-freedom omnibus test was done to test for effect in multiple reference panels. The threshold for the omnibus test was P□=□4.64E-06 (0.05/10,323 [number of genes tested]). Splicing analysis was done using the CMC dataset obtained from FUSION and an empirical p-value <0.05 were considered significant.

### Fine-mapping of TWAS associations

To address the issue of co-regulation and LD, we used FOCUS (Fine-mapping of causal gene sets) to model predicted expression correlations and to assign a posterior probability for causality in relevant tissue types^12^. Briefly, FOCUS prioritizes genes for each TWAS hit to be included in a 90%-credible set while accounting for pleiotropic SNP effects. The identical TWAS reference panels for FUSION were used as in the analysis described above.

### Phenome-wide association studies

To identify phenotypes associated with *FLT3*, a phenome-wide association study (pheWAS) was done. PheWAS was done using public data provided by GWASAtlas (https://atlas.ctglab.nl).

### Lymphoblastoid cell lines

Lymphoblastoid cell lines were prepared from consenting individuals. The biobanking of these cell lines has been approved by the institutional review board of McGill University. Cells derived from Tourette’s Syndrome patients were grown at 37°C and cells were cultured for approximately one week prior to RNA extraction.

### RNA extraction

RNA was extracted from the cells using the Qiagen RNAeasy Mini Kit. The RNA was subsequently stored in -80°C after elution with RNAse-free water. One microgram of each sample of RNA was converted into cDNA using the SuperScript VILO cDNA Synthesis Kit by Thermo Fisher Scientific.

### Reverse-transcriptase quantitative qPCR

The cDNA was used to perform a Taqman qPCR using QuantStudio 7. The *FLT3* probe was used, and *POLR2A* (polymerase [RNA] II [DNA-directed] polypeptide) was used as the endogenous control. The qPCR was performed in triplicate. A Shapiro-Wilk Test was done to determine the normality of the mean CT values data. Mean CT values were derived from averaging the triplicate CT values. An ANOVA was done using the model mean CT values ∼ sex + plate + disease status + disease status:sex). Mean CT values were derived from averaging the triplicate CT values. Cohen’s F was used to determine the effect size of the data.

## Data availability

Summary statistics have been attached as Supplementary Data 1. All other data are contained within the article or its supplementary information and upon reasonable request.

## Competing interests

The authors declare no competing interests.

## Contributions

C.L. performed analyses and drafted the manuscript. V.V. and H.C. helped with molecular work. F.A., I.A.M and J.P.R. provided scientific input. P.A.D. and G.A.R. oversaw the manuscript.

## Acknowledgements

We would like to acknowledge the Psychiatric Genetics Consortium for the aggregation and release of summary statistics. We would like to thank any patients for participating in our studies. C.L. would like to dedicate this paper to Emily B. for inspiring him to study Tourette’s Syndrome.

## Funding source

Canadian Institutes of Health Research

## Notes

**Conflict of Interests:** All authors report no conflicts of interest.

